# Genome-Resolved Characterization of Stage-Associated Microbial Composition and Functional Potential During Spontaneous Ugba Fermentation

**DOI:** 10.64898/2026.09.26.754590

**Authors:** Kelechi Stanley Dike, Ojukwu Chibueze Chukwuka, Fredrick Temitope Oladele, Ezeh Chidinma Francisca, Michael Ikechukwu Nwachukwu

**Author notes:** Corresponding author: Kelechi Stanley Dike; Kelechi Stanley Dike 0000-0002-1723-2436.

## Abstract

Ugba, a traditional alkaline-fermented condiment produced from African oil bean seeds (*Pentaclethra macrophylla* Benth.), remains poorly characterised at genome resolution. This study characterised a single spontaneous ugba fermentation using genome-resolved shotgun metagenomics across Early (0–24 h), Mid (48–96 h), and Late (120– 144 h) composite stages. Thirty metagenome-assembled genomes (MAGs) were recovered and grouped according to their temporal abundance patterns. The Early-stage retained MAG population was dominated by Bacteroidota-associated MAGs, which exhibited the highest carbohydrate-active enzyme density and broad glycoside hydrolase repertoires, consistent with substantial genome-encoded potential for utilisation of the plant-derived seed matrix. The Mid stage showed marked restructuring of MAG composition, while Core and Late specialist MAGs increased towards fermentation maturation. Functional differentiation was also evident among *Corynebacterium* MAGs: *C. nuruki* encoded a complete urease system, whereas *C. phoceense_A* encoded glutamate dehydrogenase and nitrate-reduction genes. These stage-associated changes coincided with progressive alkalinisation from pH 7.15 to 8.01. Together, the results show that temporal restructuring of reconstructed microbial populations was accompanied by differentiation in genome-encoded functional potential. This study provides a genome-resolved characterisation of microbial composition and functional potential during spontaneous ugba fermentation and expands current understanding of the microbial organisation of this traditional African fermented food.

## 1. Introduction

The spontaneous solid-state fermentation of African oil bean seeds (*Pentaclethra macrophylla* Benth.) into ugba is a traditional alkaline fermentation involving extensive microbial and biochemical transformation of the seed substrate. During processing, the firm cotyledons are converted into a softer, flavourful condiment through protein hydrolysis, reduction of antinutritional factors and accumulation of free amino acids, which collectively influence the nutritional and sensory properties of the product (Obeta, 1983; Olasupo et al., 2016). Ugba is an important traditional food in Nigeria and neighbouring regions and contributes to established local food-processing practices and economies (Odunfa, 1985; Achi, 2005; Nwokeleme and Ugwuanyi, 2015). Understanding the microorganisms and functional capacities associated with these transformations is relevant to efforts aimed at improving fermentation consistency and informing future starter-culture development.

Ugba fermentation involves degradation of complex carbohydrates, extensive protein transformation, lipid metabolism and progressive alkalinisation, reflecting the collective activities of microorganisms within a changing fermentation environment. Early microbiological studies relied mainly on culture-dependent approaches and consistently identified *Bacillus* spp. as important fermentative organisms, with lactic acid bacteria and Gram-negative taxa also reported at different stages (Obeta, 1983; Odunfa and Oyeyiola, 1985; Isu and Ofuya, 2000). These studies established the microbiological foundation of ugba fermentation but, by design, primarily characterised organisms recoverable under the cultivation conditions employed. Consequently, they provided limited information on the genome-resolved composition and encoded functional capacities of the broader microbial community.

Culture-independent approaches subsequently expanded understanding of microbial diversity in African fermented foods by detecting taxa that may be poorly represented in cultivation-based surveys (Ercolini, 2013; Adedeji et al., 2017; Diaz et al., 2019).

However, marker-gene sequencing provides limited resolution for reconstructing population genomes and for assigning metabolic capacities directly to individual microbial populations (Quince et al., 2017; De Filippis et al., 2017; Knight et al., 2018). Consequently, while such approaches are valuable for describing community composition, they provide less direct information on how genome-encoded functions are distributed among microbial populations during fermentation.

More recently, a shotgun metagenomic survey of 91 African fermented foods, including ugba, demonstrated the value of genome-resolved approaches for characterising food-associated microbial populations and their carbohydrate-active enzyme, virulence-factor and antimicrobial-resistance gene profiles (Leech et al., 2026). Because that study was designed as a broad cross-sectional survey, it did not address stage-associated changes within an individual ugba fermentation or examine how genome-encoded functional potential varied across successive fermentation stages. Thus, temporal genome-resolved information for ugba remains limited.

Several aspects of the genome-resolved organisation of ugba fermentation therefore remain poorly characterised. In particular, little is known about which microbial populations carry carbohydrate-active enzyme repertoires during early fermentation, how nitrogen-metabolism capacities are distributed among populations observed during alkaline development, and whether populations associated with different fermentation stages differ in their broader encoded functional profiles. Addressing these questions requires an approach capable of linking temporal abundance patterns with genomic functional potential at the population level.

Genome-resolved metagenomics combines shotgun sequencing with metagenome-assembled genome reconstruction to recover individual microbial populations and associate their abundance patterns with genome-encoded functional potential (Bowers et al., 2017; Srinivas et al., 2022; Walsh et al., 2023). Integration of MAG reconstruction with KEGG pathway annotation, carbohydrate-active enzyme profiling and Clusters of Orthologous Groups analysis enables population-level comparison of metabolic potential. Applied across sequential fermentation stages, this framework can reveal whether changes in reconstructed community composition are accompanied by corresponding changes in encoded functional capacities.

Accordingly, this study applied genome-resolved shotgun metagenomics, supported by physicochemical measurements, to characterise stage-associated microbial and functional dynamics within a single spontaneous ugba fermentation. Metagenome-assembled genomes were reconstructed and grouped according to temporal abundance patterns, while KEGG pathway annotation, carbohydrate-active enzyme profiling and COG analysis were used to examine differences in genome-encoded functional potential among the recovered populations. By integrating temporal abundance with genome-level functional characterisation, the study provides a detailed genome-resolved view of microbial and functional restructuring within the fermentation examined and establishes a basis for assessing the reproducibility and biotechnological relevance of these patterns in future independent fermentations.

## 2 Materials and methods

### 2.1 Sample collection and fermentation process

African oil bean seeds (*Pentaclethra macrophylla* Benth.) were purchased from a retail market in Owerri, Imo State, Nigeria, and processed into ugba using a traditional fermentation method with minor modifications (Odunfa & Oyeyiola, 1985). The seeds were washed, boiled for approximately 6 h until softened, manually dehulled, and sliced into thin cotyledon strips. The sliced cotyledons were rinsed with sterile distilled water, packaged in sterile food-grade polyethylene bags, and allowed to ferment spontaneously under ambient laboratory conditions without the addition of starter cultures.

A single spontaneous fermentation batch was analysed to characterise stage-associated microbial and functional dynamics during the fermentation process. Three stage-representative composite samples were generated, corresponding to the Early (0–24 h), Mid (48–96 h), and Late (120–144 h) fermentation stages. Each composite sample was prepared by pooling equal amounts of fermenting material collected at the designated time points within that stage. Approximately 50 g of each composite sample was aseptically transferred to sterile containers, transported on ice, and processed immediately for physicochemical analysis and shotgun metagenomic sequencing.

### 2.2 Physicochemical analysis

Physicochemical parameters were determined from composite samples representing the Early (0–24 h), Mid (48–96 h), and Late (120–144 h) fermentation stages. Each stage-representative composite sample was analysed in technical triplicate. pH was determined potentiometrically using a calibrated pH meter. Fermentation temperature was measured in situ using a calibrated digital thermometer inserted directly into the fermenting mass.

Moisture content was determined gravimetrically using the following equation:

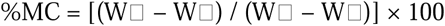

where W represents the weight of the empty crucible, W the weight of the crucible plus sample before drying, and W the weight of the crucible plus sample after oven drying. Values reported for each fermentation stage represent the mean of technical triplicate determinations. These technical replicates were used to assess analytical consistency and do not represent independent biological fermentation replicates.

### 2.3 DNA extraction and shotgun metagenomic sequencing

Total community DNA was extracted from each of the three stage-representative composite samples using the ZymoBIOMICS DNA Miniprep Kit (Zymo Research, Irvine, CA, USA) according to the manufacturer’s instructions. DNA integrity was assessed by electrophoresis on a 1% (w/v) agarose gel, while DNA concentration and purity were evaluated using a NanoDrop spectrophotometer (Thermo Fisher Scientific, Waltham, MA, USA). Sequencing libraries were prepared for paired-end shotgun metagenomic sequencing (2 × 150 bp) on the DNBSEQ-T7 platform (BGI Genomics, Shenzhen, China).

### 2.4 Quality control and metagenome assembly

Shotgun metagenomic datasets generated from the Early, Mid, and Late composite fermentation-stage samples were quality filtered and adapter trimmed using fastp v0.23.4 (Chen et al., 2018) with default paired-end parameters. High-quality reads from all three stages were co-assembled using MEGAHIT v1.2.9 (Li et al., 2015) with default metagenomic settings. Assembly quality was evaluated using MetaQUAST (Mikheenko et al., 2016).

Reads from each fermentation stage were independently mapped to the co-assembled contigs using Bowtie2 v2.5.4 (Langmead & Salzberg, 2012). The resulting alignment files were processed using SAMtools (Li et al., 2009) to generate contig coverage profiles for downstream genome binning.

### 2.5 Genome binning, refinement, and quality assessment

Metagenome-assembled genomes (MAGs) were reconstructed using a consensus binning strategy. Contig coverage profiles were generated using the jgi_summarize_bam_contig_depths script implemented in MetaBAT2 v2.12.1 (Kang et al., 2019). Genome binning was performed independently using MetaBAT2 v2.12.1, MaxBin2 v2.2.7 (Wu et al., 2016), and CONCOCT v1.1.0 (Alneberg et al., 2014). The resulting genome bins were integrated using DAS Tool (Sieber et al., 2018) to generate a consensus set of MAGs.

MAG quality was evaluated using CheckM2 v1.1.0 (Chklovski et al., 2023). Recovered genomes were classified according to the Minimum Information about a Metagenome-Assembled Genome (MIMAG) standards (Bowers et al., 2017) as high-quality MAGs (completeness ≥90% and contamination <5%) or medium-quality MAGs (completeness ≥50% and contamination <10%).

### 2.6 Taxonomic classification

Taxonomic classification of recovered MAGs was performed using GTDB-Tk v2.6.1 (Chaumeil et al., 2022) with the GTDB release r226 reference database. Species-level assignments were further evaluated using average nucleotide identity (ANI), with thresholds of ≥95% ANI and ≥20% alignment fraction applied for species-level classification.

### 2.7 MAG abundance quantification and temporal association

Relative abundances of recovered MAGs across the Early, Mid, and Late fermentation stages were quantified using CoverM v0.4.0 in genome mode. MAGs were assigned to temporal abundance groups based on their distribution across the three fermentation stages.

MAGs for which ≥70% of cumulative relative abundance occurred within a single fermentation stage were classified as Early, Mid or Late MAGs, depending on the stage of maximum representation. MAGs detected across multiple stages without marked stage dominance were classified as Transitional MAGs. MAGs consistently detected across all three stages at a relative abundance of ≥0.1% were classified as Core MAGs.

These classifications describe temporal abundance patterns within the fermentation examined. The designation “Core MAGs” refers specifically to MAGs consistently detected across all three analysed fermentation stages and does not imply that these populations constitute a universal core microbiome across independent ugba fermentations. The specialist classifications are operational designations based on temporal abundance patterns within the analysed fermentation.

### 2.8 Functional annotation and statistical analysis

Protein-coding genes were predicted using Prodigal v2.6.3 (Hyatt et al., 2010) in metagenomic mode. Predicted proteins were functionally annotated against the Clusters of Orthologous Groups (COG) database using eggNOG-mapper v2.1.13 (Cantalapiedra et al., 2021) with the eggNOG 5.0 database. KEGG Orthologs (KO) assignments generated by eggNOG-mapper were used for metabolic pathway reconstruction and comparative functional analyses.

To investigate functional differentiation among *Corynebacterium* populations associated with fermentation maturation, KEGG pathway analyses focused on *Corynebacterium nuruki* and *Corynebacterium phoceense_A*. Metabolic pathways were reconstructed by mapping KO assignments to KEGG modules and pathways (Kanehisa & Goto, 2000; Kanehisa et al., 2023).

Carbohydrate-active enzymes (CAZymes) were annotated against the CAZy database using dbCAN v5.2.8 (Drula et al., 2022), integrating HMMER3, DIAMOND, and dbCAN-sub predictions. Only CAZyme annotations supported by at least two analytical methods were retained.

Statistical enrichment of COG categories among temporal MAG groups was evaluated using Fisher’s exact test with Bonferroni correction (α = 0.05). Principal component analysis (PCA) was used as an exploratory ordination to visualise similarities and differences in COG functional profiles among individual MAGs and temporal groups. Owing to its incomplete genome recovery (54.6% completeness), bin.61 (*Kurthia gibsonii*) was retained for abundance profiling but excluded from the COG and CAZyme analyses. All statistical analyses and data visualisations were performed in R v4.3.0 (R Core Team, 2023).

## 3 Results

### 3.1 Physicochemical changes during ugba fermentation

The physicochemical characteristics of ugba across the Early (0–24 h), Mid (48–96 h), and Late (120–144 h) fermentation stages are presented in Table 1. The pH increased progressively from 7.15 at the Early stage to 8.01 at the Late stage, indicating gradual alkalinisation of the fermenting substrate. Moisture content increased modestly from 51.67% at the Early stage to 53.67% at the Late stage. Fermentation temperature remained relatively stable, ranging from 32.33 to 33.05 °C across the three stages. Overall, progressive alkalinisation represented the most pronounced physicochemical change observed across the fermentation stages.

**Table 1.** Physicochemical characteristics of Ugba during spontaneous fermentation

| Fermentation stage | pH | Moisture content (%) | Temperature (°C) |
| --- | --- | --- | --- |
| Early (0–24 h) | 7.15 | 51.67 | 32.33 |
| Mid (48–96 h) | 7.62 | 52.83 | 33.05 |
| Late (120–144 h) | 8.01 | 53.67 | 33.03 |

Values represent the mean of three technical determinations performed on each stage-representative composite sample. Technical replicates were used to assess analytical consistency and do not represent independent biological fermentation replicates.

### 3.2 Metagenome assembly and MAG quality assessment

Quality-filtered reads from the Early, Mid, and Late fermentation stages were co-assembled using MEGAHIT, generating a 458.6 Mb assembly comprising 110,194 contigs (≥1,000 bp) with an N50 of 8,408 bp. Assembly-level read recruitment ranged from 89.4% to 95.8% across fermentation stages, indicating that most quality-filtered reads were represented in the co-assembly.

Genome binning using MetaBAT2, MaxBin2, CONCOCT, and DAS Tool generated 38 genome bins, of which 30 met the specified quality criteria and were retained for downstream analyses. These comprised 18 high-quality and 12 medium-quality MAGs according to the MIMAG classification applied in this study, while the remaining eight bins were excluded because of low completeness, high contamination, or both (Fig. 1; Supplementary Tables S1 and S2).

**Figure 1.**
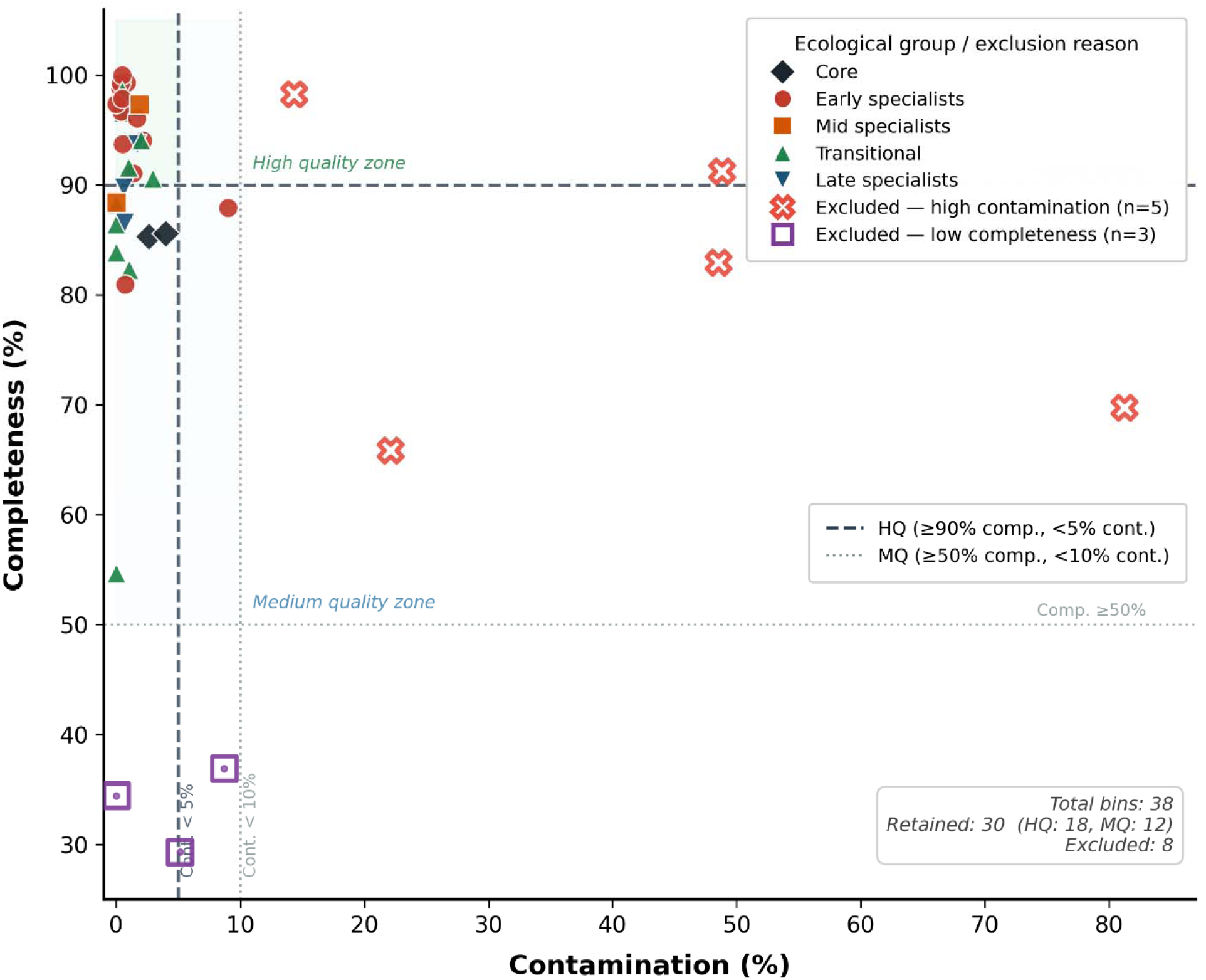
Quality assessment and selection of metagenome-assembled genomes (MAGs) recovered during spontaneous ugba fermentation. Completeness and contamination estimates of the 38 genome bins recovered from shotgun metagenomic sequencing. Retained MAGs and excluded bins are shown according to MIMAG-based quality criteria. Dashed lines indicate thresholds for high-quality and medium-quality MAG classification.

The retained MAGs exhibited completeness values ranging from 54.6% to 100.0% and contamination levels between 0.0% and 9.0%, with most genomes occupying the high-completeness, low-contamination region of the quality space (Fig. 1). *Kurthia gibsonii* (bin.61) was retained for abundance profiling but excluded from COG an CAZyme analyses because its relatively low completeness (54.6%) limited confidence in genome-resolved functional inference.

### 3.3 Taxonomic composition and temporal classification of recovered MAGs

Based on the predefined temporal abundance criteria, the 30 retained MAGs were assigned to five temporal groups: Early-associated, Mid-associated, Transitional, Late-associated, and Core MAGs (Fig. 2; Table 2). Taxonomic classification identified 26 genera distributed across four bacterial phyla: Actinomycetota (9 MAGs), Bacillota (9 MAGs), Bacteroidota (8 MAGs), and Pseudomonadota (4 MAGs). Nineteen MAGs were assigned to species level, while the remaining 11 were classified at the genus level (Table 2).

**Figure 2.**
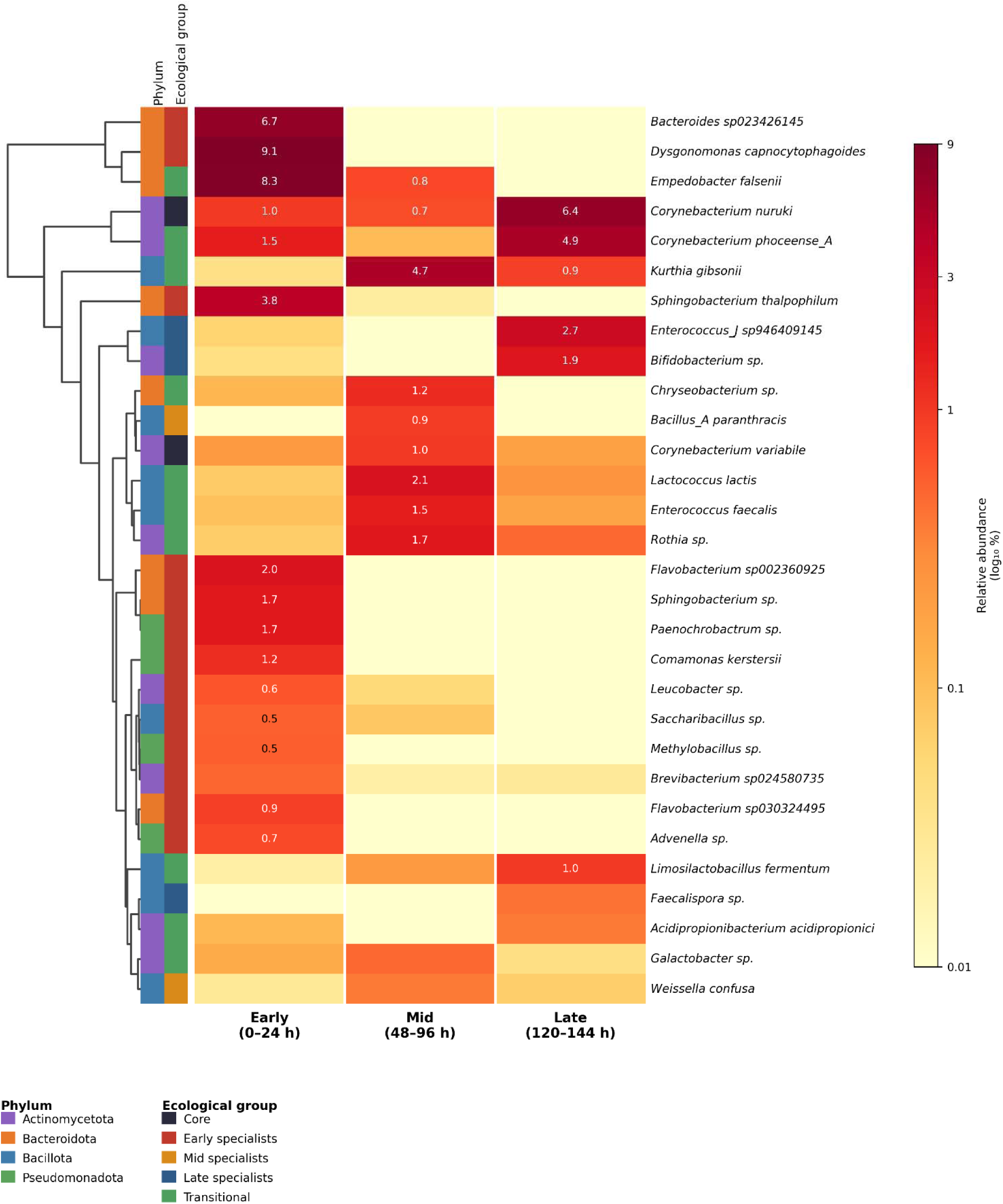
Genome-resolved temporal abundance patterns and taxonomic classification of retained MAGs during spontaneous ugba fermentation. Heatmap showing log10-transformed relative abundance of the 30 retained MAGs across Early (0–24 h), Mid (48– 96 h), and Late (120–144 h) fermentation stages. Rows represent MAGs grouped according to temporal abundance classifications, with taxonomic affiliations indicated.

**Table 2.** Taxonomic composition and ecological classification of the 30 retained metagenome-assembled genomes (MAGs) recovered during ugba fermentation.

| Temporal group | No. of MAGs | Representative taxa | Dominant phylum |
| --- | --- | --- | --- |
| Early specialists | 13 | *Dysgonomonas capnocytophagoides*,<br>*Bacteroides* sp.,<br>*Sphingobacterium* spp.,<br>*Flavobacterium* spp. | Bacteroidota (highest-abundance MAGs) |
| Mid specialists | 2 | *Bacillus_A paranthracis*;<br>*Weissella confusa* | Bacillota |
| Transitional | 10 | *Empedobacter falsenii*,<br>*Kurthia gibsonii*,<br>*Lactococcus lactis*,<br>*Corynebacterium phoceense_A* | Mixed |
| Late specialists | 3 | *Enterococcus_J* sp.,<br>*Bifidobacterium* sp.,<br>*Faecalispora* sp. | Mixed |
| Core | 2 | *Corynebacterium variabile*;<br>*Corynebacterium nuruki* | Actinomycetota |

Early-associated MAGs comprised 13 genomes and included predominantly Bacteroidota-affiliated taxa, such as *Dysgonomonas capnocytophagoides*, *Bacteroides* sp., *Sphingobacterium* spp., and *Flavobacterium* spp. The two Mid-associated MAGs, *Bacillus_A paranthracis* and *Weissella confusa*, belonged to Bacillota. Transitional MAGs (n = 10) represented the most taxonomically diverse group, encompassing members of Actinomycetota, Bacillota, Bacteroidota, and Pseudomonadota. The three Late-associated MAGs included *Enterococcus_J* sp., *Bifidobacterium* sp., and *Faecalispora* sp., whereas the two Core MAGs, *Corynebacterium variabile* and *Corynebacterium nuruki*, belonged to Actinomycetota and were detected across all three fermentation stages. The recovered MAGs showed broad phylogenetic diversity and were distributed among distinct temporal abundance groups within the fermentation examined (Table 2; Fig 2).

### 3.4 Stage-associated temporal abundance dynamics of retained MAG populations during ugba fermentation

The composition of the retained MAG population changed markedly across the three fermentation stages (Fig. 3). At the Early stage, Early-associated MAGs accounted for 71.9% of the retained MAG population, with Transitional MAGs present at lower abundance, whereas Core and Late-associated MAGs together contributed only a minor fraction. At the Mid stage, the retained MAG population shifted markedly, with Transitional MAGs accounting for 80.6% of the total, while Early-associated MAGs declined to negligible abundance. *Kurthia gibsonii* and the Mid-associated *Weissella confusa* exhibited their highest relative abundances at this stage.By the Late stage, the retaine MAG population was more evenly distributed among temporal groups. Transitional MAGs remained the largest group, while Core and Late-associated MAGs increased to 33.5% and 25.3%, respectively, together accounting for more than half of the retained MAG population.

**Figure 3.**
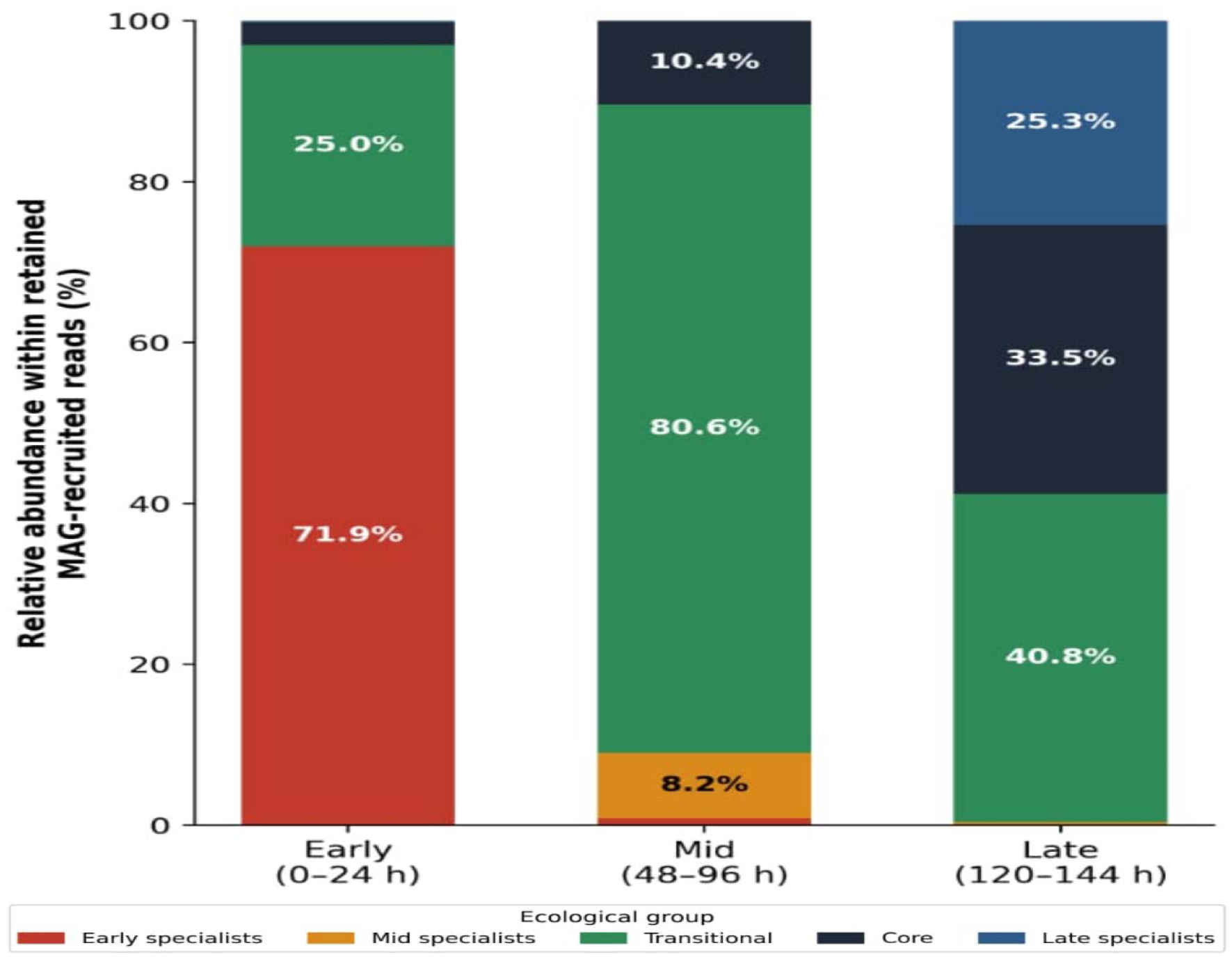
Stage-associated abundance dynamics of temporal MAG groups during spontaneous ugba fermentation. Stacked bar chart showing the relative contribution of Early-associated, Mid-associated, Transitional, Core, and Late-associated MAG groups across fermentation stages based on MAG recruitment profiles.

Overall, the fermentation examined showed marked stage-associated restructuring of the retained MAG population, progressing from Early-associated MAG predominance to a Transitional-dominated Mid stage and subsequently to increased representation of Core and Late-associated MAGs during the Late stage (Fig. 3).

### 3.5 KEGG-based functional differentiation of *Corynebacterium* MAGs associated with fermentation maturation

To examine genome-encoded functional differentiation among *Corynebacterium* populations observed during fermentation maturation, KEGG Orthology (KO)-based pathway reconstruction was performed for *Corynebacterium nuruki* (bin.5) and *Corynebacterium phoceense_A* (bin.99) (Fig. 4). A total of 1,198 and 1,118 unique KOs were identified in *C. nuruki* and *C. phoceense_A*, respectively, indicating shared metabolic capacities alongside differences in nitrogen-metabolism potential.

**Figure 4.**
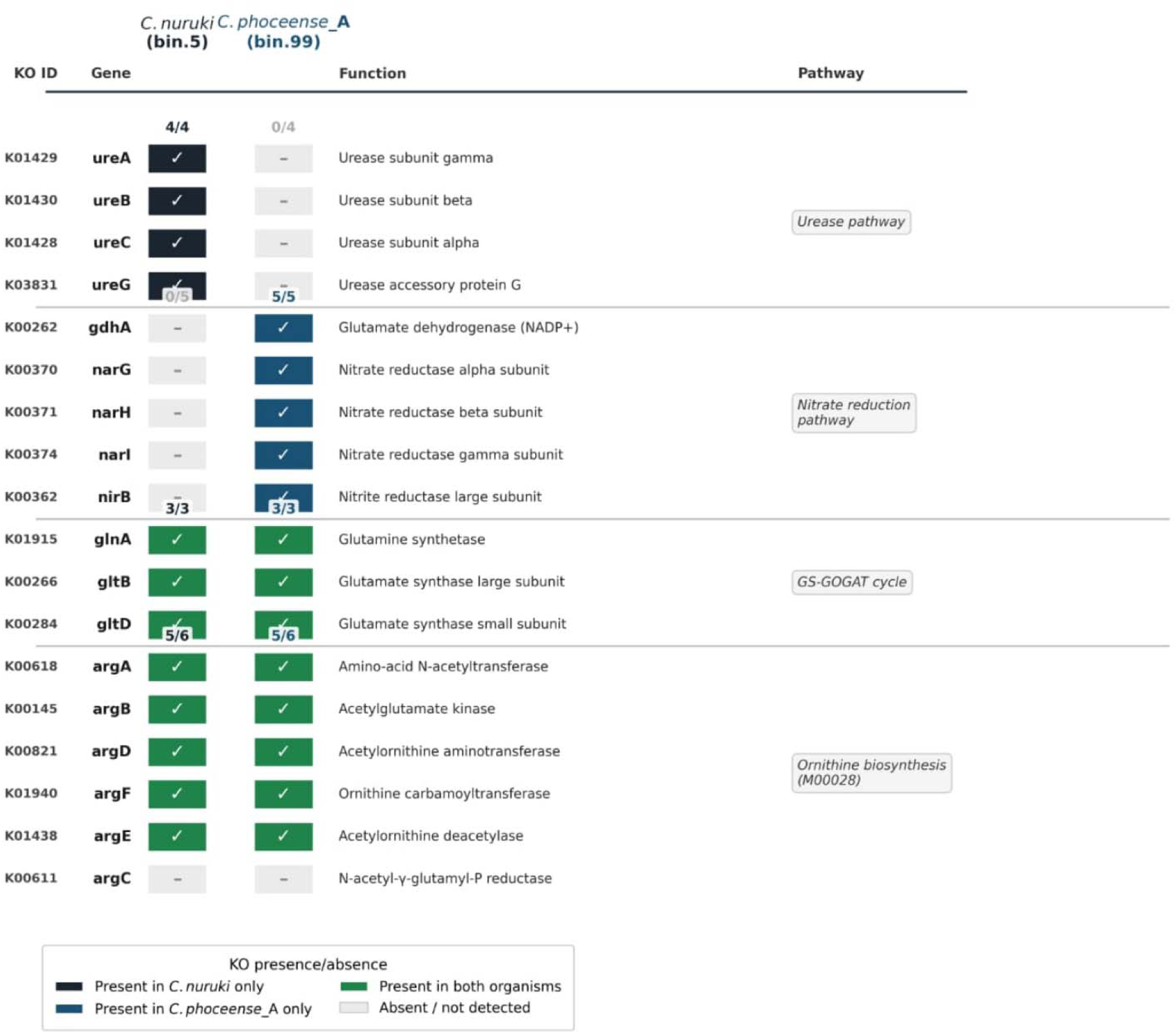
KEGG-based comparison of nitrogen-metabolism potential between Corynebacterium MAGs during ugba fermentation. Presence and absence of selected KEGG Orthology (KO) genes associated with urease, nitrate reduction, the GS– GOGAT cycle, and ornithine biosynthesis in *Corynebacterium nuruki* (bin.5) and *Corynebacterium phoceense_A* (bin.99). Coloured boxes indicate detected KOs and grey boxes indicate undetected KOs.

Both MAGs encoded a complete glutamine synthetase–glutamate synthase (GS–GOGAT) cycle and a near-complete ornithine biosynthesis pathway (5/6 KOs), indicating shared genomic potential for nitrogen assimilation and amino-acid biosynthesis (Fig. 4). However, the two MAGs differed in genes associated with ammonia-generating and nitrogen-transformation pathways.

*Corynebacterium nuruki*, which increased in relative abundance towards the Late stage, uniquely encoded a complete urease system (*ureA, ureB, ureC,* and *ureG*), whereas these genes were not detected in *C. phoceense_A*. In contrast, *C. phoceense_A* uniquely encoded glutamate dehydrogenase (*gdhA*; K00262), together with nitrate-reduction genes (*narGHI* and *nirB*), which were not detected in *C. nuruki* (Fig. 4).

These contrasting genomic configurations indicate differentiation in nitrogen-metabolism potential between the two *Corynebacterium* MAGs observed during fermentation maturation.

### 3.6 COG functional enrichment of Early-associated MAGs

Comparative COG profiling of Early-associated MAGs against all other temporal MAG groups identified 18 significantly differentiated functional categories following Fisher’s exact test with Bonferroni correction, comprising 10 enriched and eight depleted categories (Fig. 5).

**Figure 5.**
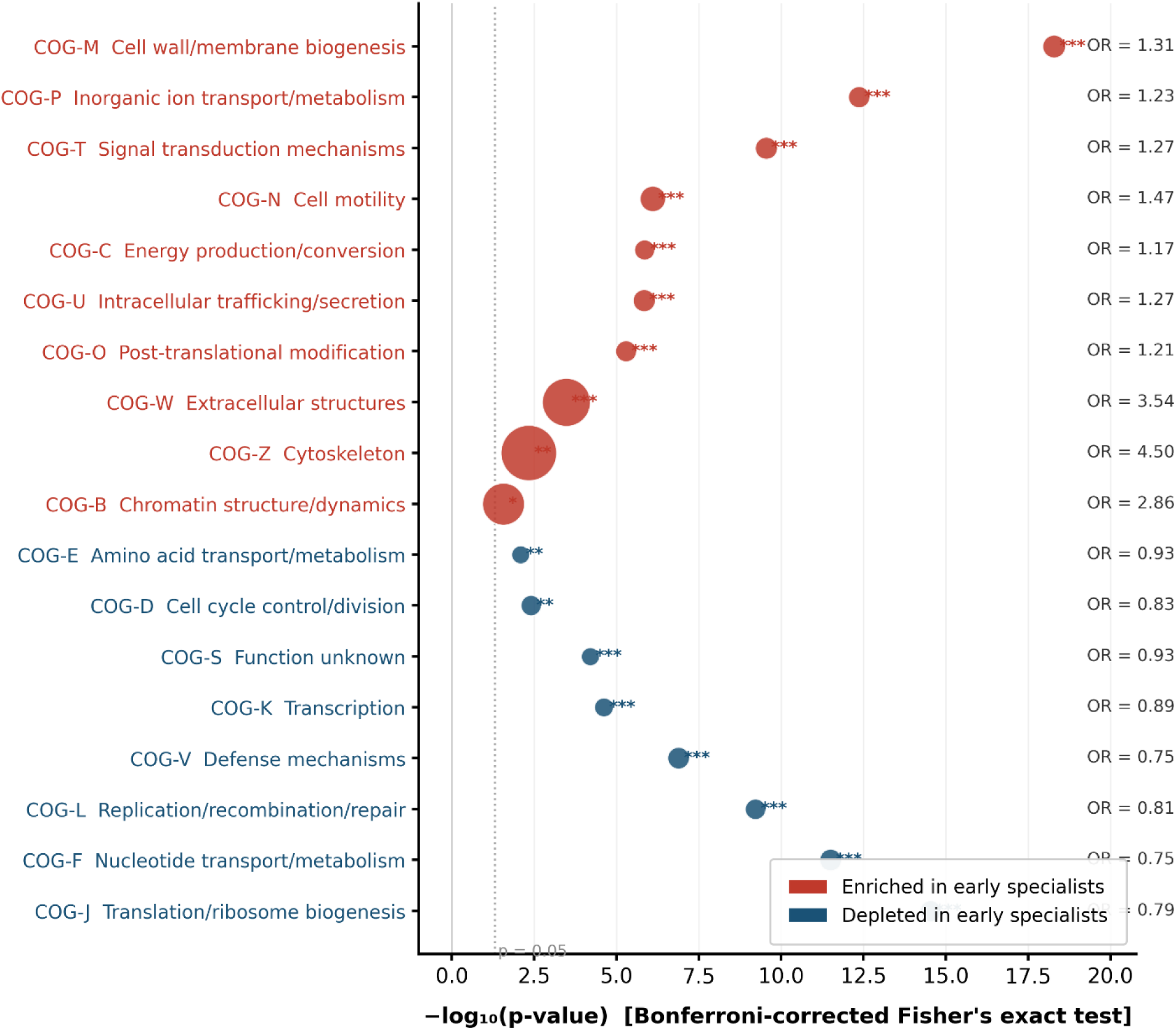
Differential COG functional enrichment among Early-associated MAGs. Bubble plot showing significantly enriched and depleted COG functional categories in Early-associated MAGs compared with other temporal MAG groups. Statistical significance was assessed using Fisher’s exact test with Bonferroni correction. Bubble size represents odds ratio, and colour indicates enrichment or depletion.

Among the enriched functions, categories associated with cell envelope biogenesis, inorganic ion transport, signal transduction, cell motility, energy production, intracellular trafficking and secretion, and post-translational modification were significantly overrepresented among Early-associated MAGs. Cell wall and membrane biogenesis (COG-M), inorganic ion transport and metabolism (COG-P), and signal transduction mechanisms (COG-T) exhibited the strongest statistical enrichment, whereas extracellular structures (COG-W), cytoskeleton (COG-Z), and chromatin structure and dynamics (COG-B) showed the largest effect sizes despite their lower gene representation (Fig. 5)

Conversely, Early-associated MAGs were significantly depleted in functional categories associated with translation, nucleotide transport and metabolism, replication and DNA repair, defence mechanisms, transcription, cell cycle control, amino-acid transport and metabolism, and genes of unknown function. Translation and ribosome biogenesis (COG-J) and nucleotide transport and metabolism (COG-F) exhibited the strongest statistical depletion (Fig. 5).

### 3.7 CAZyme class profiles across temporal MAG groups

CAZyme annotation of the 29 MAGs included in the functional analyses, excluding bin.61 because of low completeness, revealed differences in CAZyme repertoire size, density and class composition among temporal MAG groups (Fig. 6). Early-associated MAGs encoded the largest CAZyme repertoire, averaging 114.6 CAZymes per genome, and exhibited the highest CAZyme density, with 33.1 CAZymes per 1,000 genes. In contrast, Core MAGs encoded the smallest CAZyme repertoire, averaging 41.0 CAZymes per genome, and exhibited the lowest CAZyme density, with 16.3 CAZymes per 1,000 genes. Early-associated MAGs displayed a distinct genome-encoded functional profile characterised by enrichment of categories associated with environmental sensing, cell-envelope processes and resource acquisition, together with depletion of functions related to information storage, genome maintenance and core cellular metabolism (Fig. 6).The proportional distribution of major CAZyme classes also varied among temporal MAG groups. GTs and GHs constituted the largest CAZyme classes among Early-associate MAGs, whereas Late-associated MAGs exhibited the highest proportional representation of GHs (53%). Core MAGs showed the highest proportions of CEs (28%) and GTs (45%), while Mid-associated MAGs displayed the highest proportional representation of CBMs (12%). Transitional MAGs exhibited an intermediate class distribution but contained a higher proportion of CEs (18%) than the Early-, Mid-, and Late-associated groups (Fig. 6).

**Figure 6.**
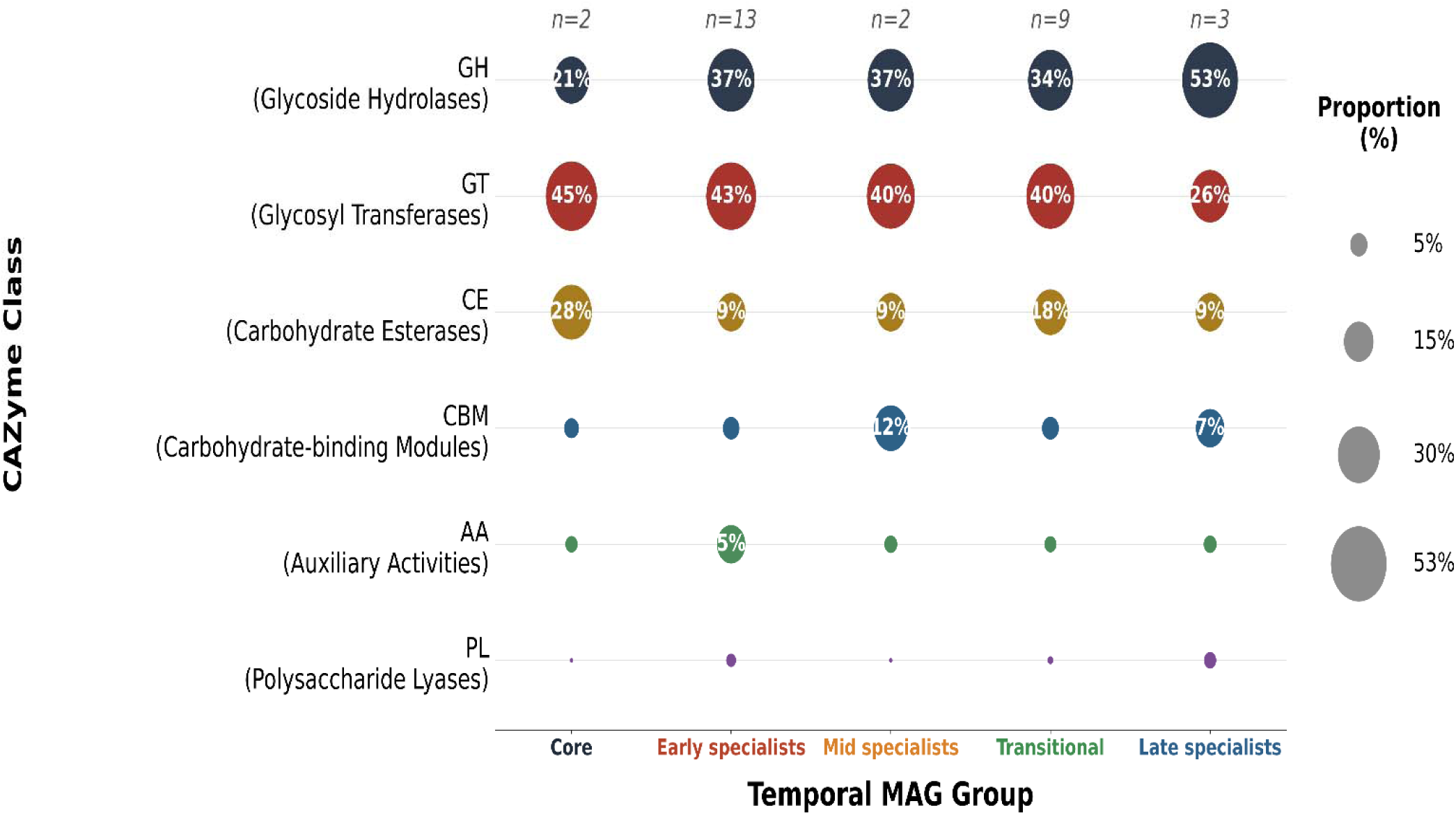
Carbohydrate-active enzyme class composition across temporal MAG groups during spontaneous ugba fermentation. Bubble plot showing the proportional distribution of CAZyme classes, including GH, GT, CE, CBM, AA, and PL, across temporal MAG groups. Analysis included 29 MAGs, excluding bin.61 (*Kurthia gibsonii*) because of low genome completeness. Group sizes are indicated above each column.

### 3.8 Glycoside hydrolase family distribution across temporal MAG groups

Analysis of glycoside hydrolase (GH) family distributions across temporal MAG groups identified 28 GH families, of which the 20 most abundant are presented in Fig. 7. GH family composition differed among groups, with Early-associated MAGs encoding the broadest and most abundant GH repertoire.

**Figure 7.**
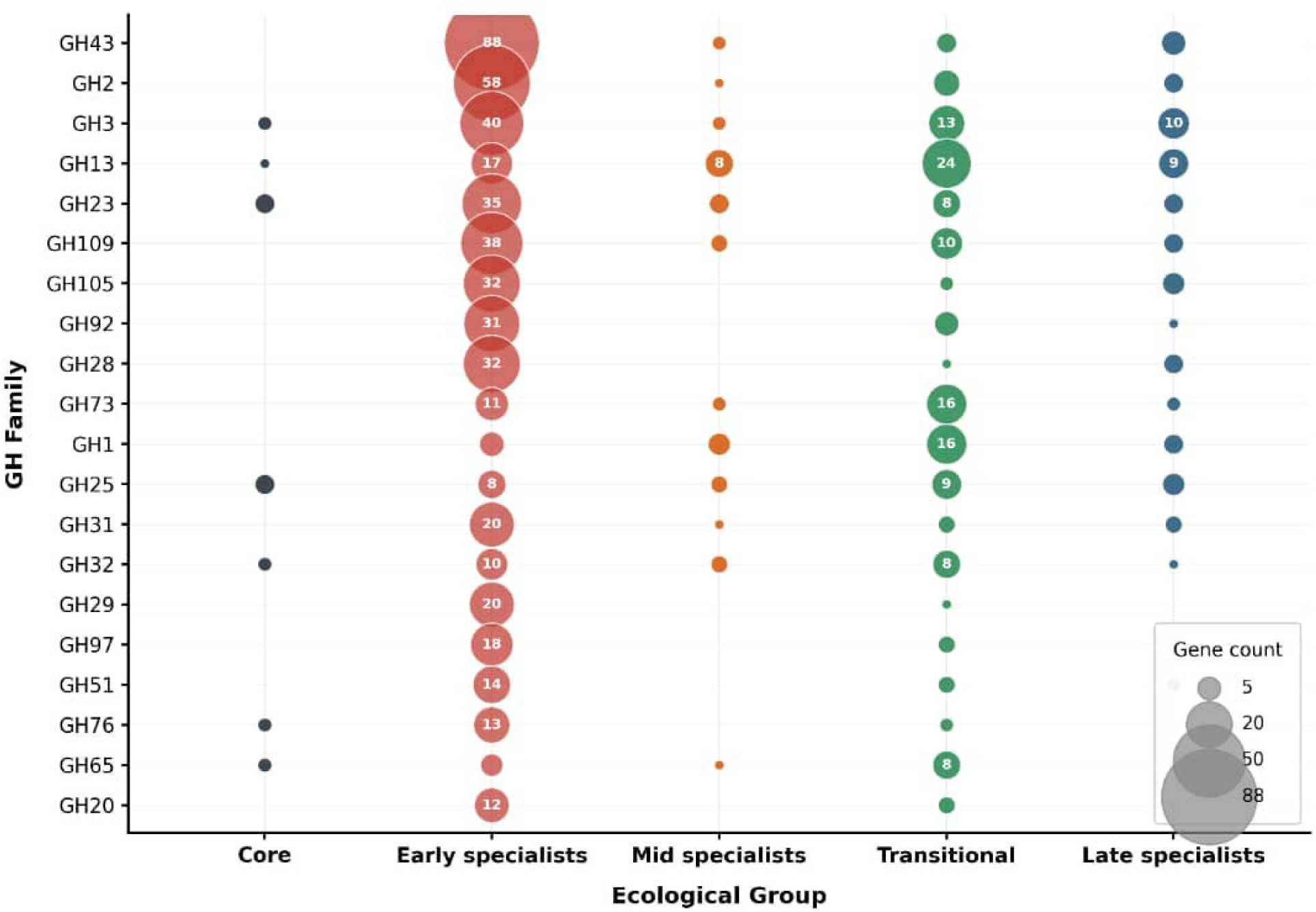
Distribution of glycoside hydrolase families across temporal MAG groups during spontaneous ugba fermentation. Bubble plot showing the distribution of the 20 most abundant glycoside hydrolase (GH) families across temporal MAG groups. Bubble size represents the number of GH genes assigned to each family, with values shown inside bubbles. Analysis included 29 MAGs after exclusion of bin.61 (*Kurthia gibsonii*).

GH43 was the most abundant GH family and was strongly concentrated among Early-associated MAGs, which encoded 88 GH43 genes. Early-associated MAGs also accounted for most genes assigned to several other highly represented families, including GH2, GH105, GH92, GH28, and GH109 (Fig. 7). In contrast, GH13 showed a broader temporal distribution and was most abundant among Transitional MAGs, which encoded 24 genes compared with 17 among Early-associated MAGs. GH73 was similarly more abundant among Transitional MAGs (16 genes) than among Early-associated MAGs (11 genes).

Late-associated MAGs exhibited a comparatively restricted GH repertoire, although several families, including GH3, GH13, GH105, GH28, GH1, GH25, and GH31, were represented. Core MAGs also showed a restricted GH repertoire, with genes detected in a limited number of families, including GH3, GH13, GH23, GH25, GH32, GH76, and GH65 (Fig. 7).

GH family distributions differed among temporal MAG groups, with Early-associated MAGs contributing strongly to most of the abundant GH families, while Transitional MAGs showed greater representation of selected families, particularly GH13 and GH73. Core and Late-associated MAGs exhibited comparatively restricted GH family profiles (Fig. 7).

### 3.9 Functional differentiation among temporal MAG groups

Principal component analysis (PCA) of COG-based functional profiles across the 29 MAGs included in the functional analyses, with bin.61 excluded because of low completeness, showed differences in the distribution of temporal MAG groups along the first two principal components, which together explained 40.9% of the total functional variance (PC1 = 21.8%; PC2 = 19.1%) (Fig. 8A). Given the unequal numbers of MAGs among temporal groups, particularly the small Mid-associated and Core groups (n = 2 each), the PCA was interpreted as a exploratory representation of genome-level functional variation rather than as evidence of statistically supported separation among groups.

**Figure 8.**
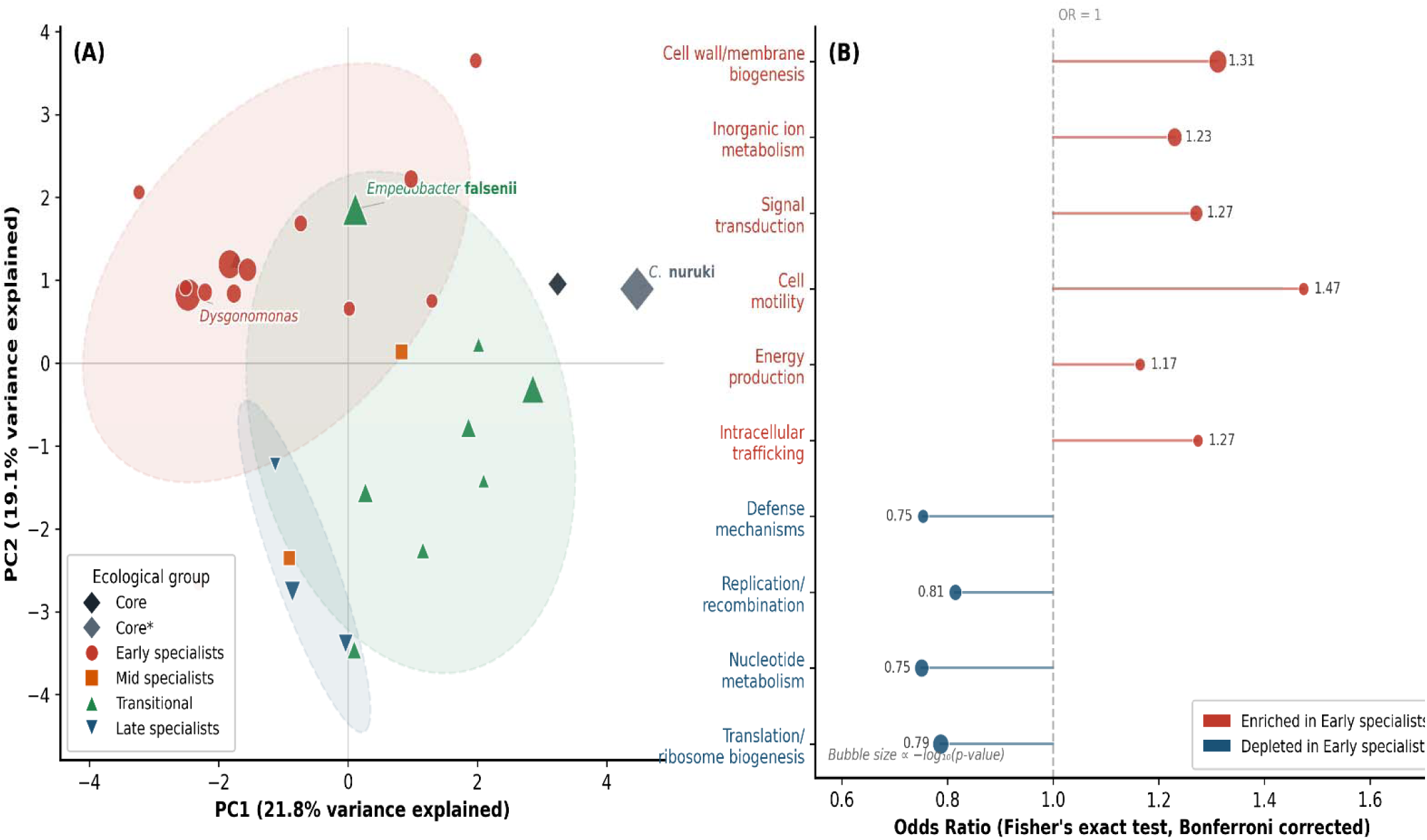
Functional differentiation among temporal MAG groups based on COG profiles. (A) PCA of COG functional profiles among MAGs included in functional analysis. Symbols represent temporal MAG groups and shaded regions indicate group dispersion. (B) Odds ratios for significantly enriched or depleted COG categories in Early-associated MAGs compared with other temporal MAG groups. Enrichment analysis was performed using Fisher’s exact test with Bonferroni correction.

Early-associated MAGs clustered predominantly towards the negative PC1 region, whereas Core MAGs occupied the positive PC1 region of the ordination space. Transitional MAGs were more broadly dispersed, spanning the central and positive PC1 regions, while the two Mid-associated MAGs occupied separate positions within the ordination. Late-associated MAGs were positioned primarily within the lower portion of the ordination space (Fig. 8A).

The distribution of temporal MAG groups in the PCA was consistent with the functional differences identified by COG enrichment analysis. Early-associated MAGs were enriched in categories including cell wall and membrane biogenesis, inorganic ion transport and metabolism, signal transduction mechanisms, cell motility, energy production, and intracellular trafficking, while categories associated with defence mechanisms, replication and recombination, nucleotide metabolism, and translation and ribosome biogenesis were depleted (Fig. 8B).

Together, the PCA ordination and COG enrichment analysis indicate functional differentiation among temporal MAG groups and are consistent with stage-associated redistribution of genome-encoded functional potential within the fermentation examined (Fig. 8).

## 4. Discussion

The spontaneous ugba fermentation examined in this study showed marked stage-associated restructuring of genome-resolved microbial populations, accompanied by progressive alkalinisation and differences in genome-encoded functional potential among temporal MAG groups. The Early stage was characterised by a high representation of Bacteroidota-associated MAGs with extensive carbohydrate-active enzyme repertoires, followed by a Transitional-dominated Mid stage and increased representation of Core and Late-associated MAGs towards fermentation maturation. Taken together, these observations indicate that changes in the composition of the reconstructed microbial population within this fermentation were accompanied by redistribution of encoded functional capacities.

A notable observation was the high representation of Bacteroidota-associated MAGs during the Early stage, contrasting with the historical emphasis on *Bacillus* spp. in culture-based studies of ugba fermentation (Obeta, 1983; Odunfa & Oyeyiola, 1985). Prominent Early-associated populations included *Dysgonomonas*, *Bacteroides*, *Sphingobacterium* and *Flavobacterium*. These MAGs exhibited the highest CAZyme density and broad glycoside hydrolase repertoires among the temporal groups, indicating substantial genome-encoded capacity for utilisation of plant-derived polysaccharides. Their enrichment in COG categories associated with cell-envelope processes, signal transduction, motility and inorganic-ion transport further suggests genomic traits compatible with environmental responsiveness and resource acquisition. These features are consistent with the recognised carbohydrate-degrading capacities of several members of Bacteroidota (Grondin et al., 2017). However, because the present analysis characterises genomic potential rather than measured metabolic activity, the specific contribution of these populations to seed-matrix degradation requires experimental validation.

The transition from the Early to the Mid stage was accompanied by marked compositional turnover within the retained MAG population. Early-associated MAGs declined substantially, while Transitional MAGs became predominant. *Kurthia gibsonii*, classified as a Transitional MAG, reached its highest relative abundance during the Mid stage. The occurrence of *K. gibsonii* in ugba has recently been reported using 16S rRNA sequence analysis (Nwaiwu et al., 2026), although its temporal abundance pattern and genome-level characteristics were not resolved. The present observations therefore add genome-resolved temporal information for this population within the fermentation examined.

Fermentation maturation was accompanied by increased representation of *Corynebacterium* populations, particularly *Corynebacterium nuruki*. This species increased in relative abundance towards the Late stage, which also corresponded with the highest pH recorded during the fermentation. *C. nuruki* was originally isolated from nuruk, a Korean traditional alcohol-fermentation starter (Shin et al., 2011), making its detection and increasing representation in a fermented plant substrate noteworthy.

Genome-level comparison further revealed contrasting nitrogen-metabolism potential between *C. nuruki* and *C. phoceense_A*. *C. nuruki* encoded a complete urease system, whereas *C. phoceense_A* encoded glutamate dehydrogenase (*gdhA*) together with nitrate-reduction genes (*narGHI* and *nirB*). These contrasting genomic configurations indicate differentiation in nitrogen-acquisition and transformation potential between the two populations. Urease can generate ammonia through urea hydrolysis (Mobley & Hausinger, 1989), while glutamate dehydrogenase and nitrate-reduction pathways participate in broader microbial nitrogen metabolism (Moreno-Vivián et al., 1999; Reitzer, 2003). The temporal increase in *Corynebacterium* MAGs together with progressive alkalinisation therefore raises the possibility that nitrogen-transforming populations may contribute to late-stage alkaline development. Nevertheless, the present data demonstrate genomic potential and temporal association rather than pathway activity or causality, and direct validation using metabolomic, transcriptomic or culture-based functional approaches would be required to establish the contribution of these populations to ammonia production and pH change.

Differences in CAZyme composition, glycoside hydrolase repertoires and COG functional profiles further showed that temporal MAG groups differed in their encoded functional capacities. Early-associated MAGs were characterised by comparatively extensive carbohydrate-active enzyme repertoires and COG profiles associated with environmental sensing and resource acquisition, whereas Transitional, Core and Late-associated MAGs occupied differentiated functional space in the PCA. These patterns are consistent with stage-associated redistribution of genome-encoded functional potential within the fermentation examined rather than taxonomic restructuring alone.

The genome-resolved patterns observed here complement the predominantly culture-based understanding of ugba microbiology. Culture-dependent studies have consistently identified *Bacillus* spp. as important organisms associated with ugba fermentation (Obeta, 1983; Odunfa & Oyeyiola, 1985; Nwanekwu & Dike, 2018; Olasupo et al., 2016), while more recent molecular approaches have expanded the recognised diversity of the associated microbiota (Nwaiwu et al., 2026). In the present metagenomic dataset, *Bacillus*-affiliated MAGs did not represent the dominant fraction of the reconstructed MAG population at the analysed stages. This difference should not be interpreted as contradicting culture-based findings, because cultivation and metagenomic sequencing interrogate different components of microbial communities. Culture-based methods characterise viable organisms recoverable under defined growth conditions, whereas shotgun metagenomics provides a broader DNA-based representation and permits reconstruction of genome-encoded functional potential. Integration of both approaches in future replicated studies would provide a more complete understanding of viable, abundant and functionally relevant populations during ugba fermentation.

The single-fermentation design represents an important limitation of the present study. The temporal patterns described here therefore apply to the fermentation examined and should not be interpreted as establishing a reproducible population-level succession model for ugba fermentation. Independent biological fermentations conducted across processing conditions, seasons and locations will be required to determine the reproducibility of the observed stage-associated patterns. Nevertheless, the convergence of genome-resolved taxonomic profiles, temporal abundance patterns, functional annotations and physicochemical measurements provides a detailed characterisation of microbial and functional restructuring within this fermentation. Future studies integrating replicated fermentations with metatranscriptomic, metabolomic or targeted functional analyses would be particularly valuable for determining whether the encoded functional capacities identified here are consistently expressed and contribute directly to substrate transformation and alkaline development (Ferrocino et al., 2023).

## 5. Conclusion

This study provides a genome-resolved characterisation of stage-associated microbial and functional dynamics within a single spontaneous ugba fermentation. The Early stage was characterised by a high representation of Bacteroidota-associated MAGs with extensive carbohydrate-active enzyme and glycoside hydrolase repertoires, followed by marked restructuring of the retained MAG population during the Mid stage and increased representation of Core and Late-associated MAGs towards fermentation maturation.

Genome-level functional analyses further revealed differences in CAZyme and COG profiles among temporal MAG groups and contrasting nitrogen-metabolism potential between *Corynebacterium nuruki* and *Corynebacterium phoceense_A*. Within the fermentation examined, these stage-associated changes in reconstructed microbial populations were accompanied by redistribution of genome-encoded functional potential.

The findings provide a detailed genome-resolved framework for understanding microbial and functional organisation during ugba fermentation and identify microbial populations and encoded capacities that warrant further investigation. Because the study was based on a single fermentation batch, independent biological fermentations will be required to determine the reproducibility of the observed temporal patterns and to establish the contribution of the identified genomic functions to substrate transformation and alkaline development.

## Supplementary information

Supplementary material associated with this article is available online Statements & Declarations

## Competing Interests

The authors declare no competing interests. Author Contributions (CRediT)

Kelechi Stanley Dike: Conceptualisation, Methodology, Investigation, Formal analysis, Writing – original draft, Writing – review & editing.

Ojukwu Chibueze Chukwuka: Investigation, Formal analysis, Resources

Fredrick Temitope Oladele: Data curation, Software, Formal analysis, Writing – review & editing. Ezeh Chidinma Francisca: Methodology, Writing – review & editing.

Michael Ikechukwu Nwachukwu: Supervision

## Data Availability Statement

The metagenome-assembled genomes (MAGs) generated during this study have been deposited in the National Centre for Biotechnology Information (NCBI) under BioProject accession number PRJNA1499312. Individual BioSample accession numbers and genome quality metrics for the metagenome-assembled genomes (MAGs) are provided in Supplementary Table S4.

## Supplementary Material

**Supplementary Table S1.**
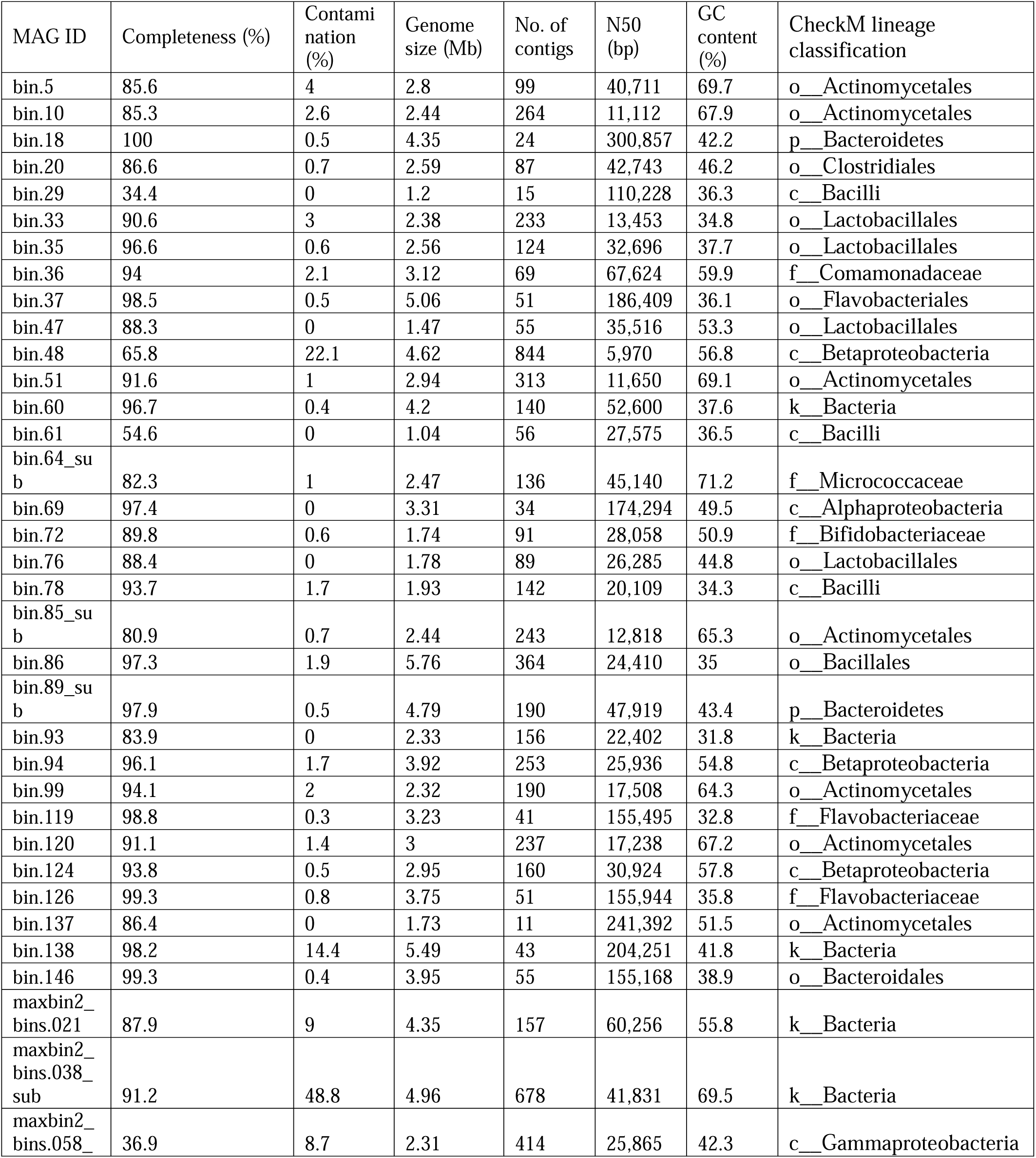
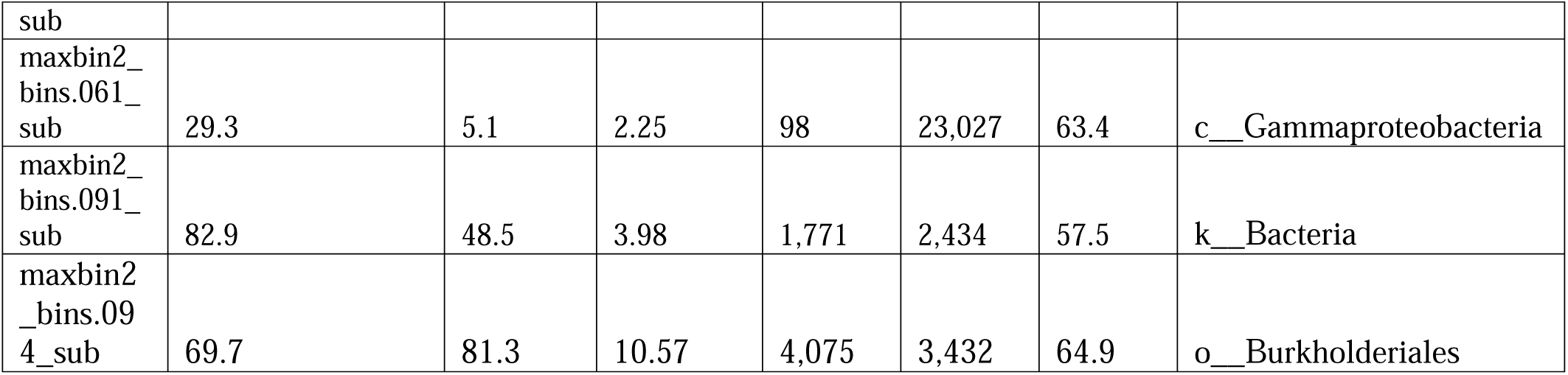
Quality statistics and assembly metrics of retained metagenome-assembled genomes (MAGs) recovered during ugba fermentation.

**Supplementary Table S2.** Excluded MAGs and reasons for exclusion.

| Bin ID | Completeness (%) | Contamination (%) | Reason for exclusion |
| --- | --- | --- | --- |
| bin.29 | 34.4 | 0 | Insufficient completeness |
| bin.48 | 65.8 | 22.1 | Excessive contamination |
| bin.138 | 98.2 | 14.4 | Excessive contamination |
| maxbin2_bins.038_sub | 91.2 | 48.8 | Excessive contamination |
| maxbin2_bins.058_sub | 36.9 | 8.7 | Insufficient completeness |
| maxbin2_bins.061_sub | 29.3 | 5.1 | Insufficient completeness |
| maxbin2_bins.091_sub | 82.9 | 48.5 | Excessive contamination |
| maxbin2_bins.094_sub | 69.7 | 81.3 | Excessive contamination |

**Supplementary Table 3.**
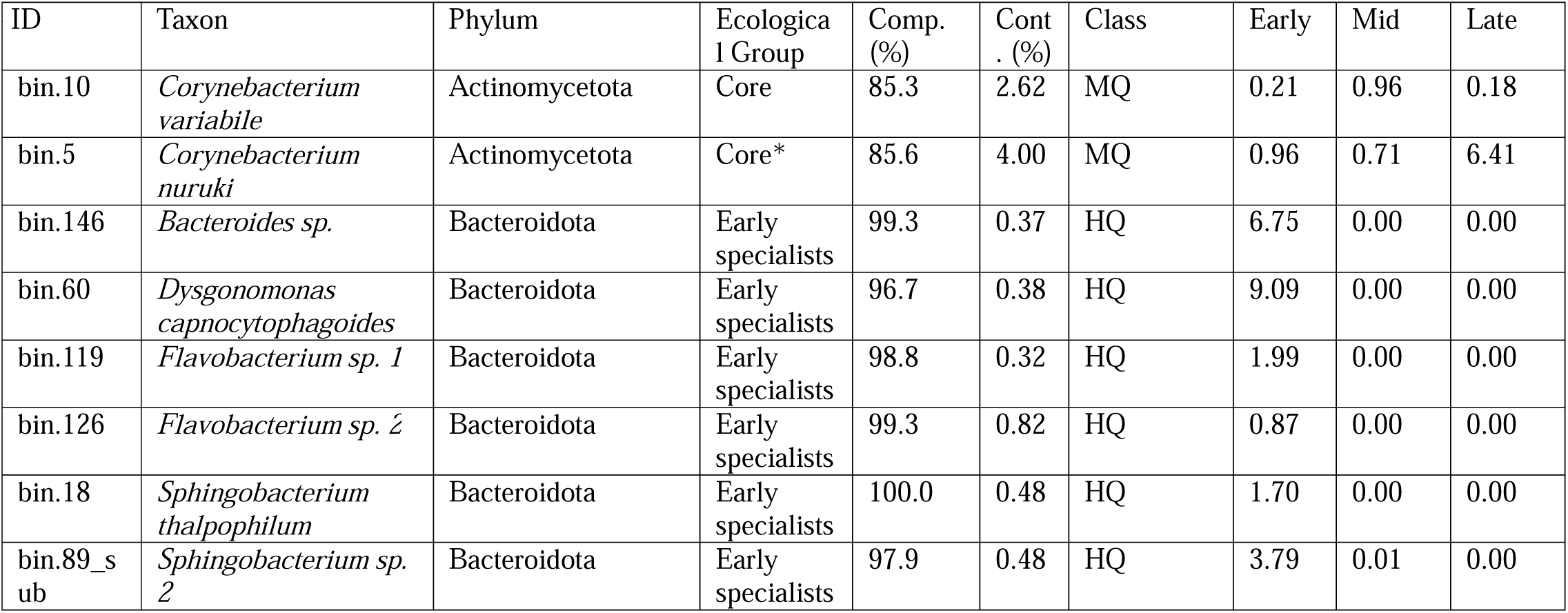
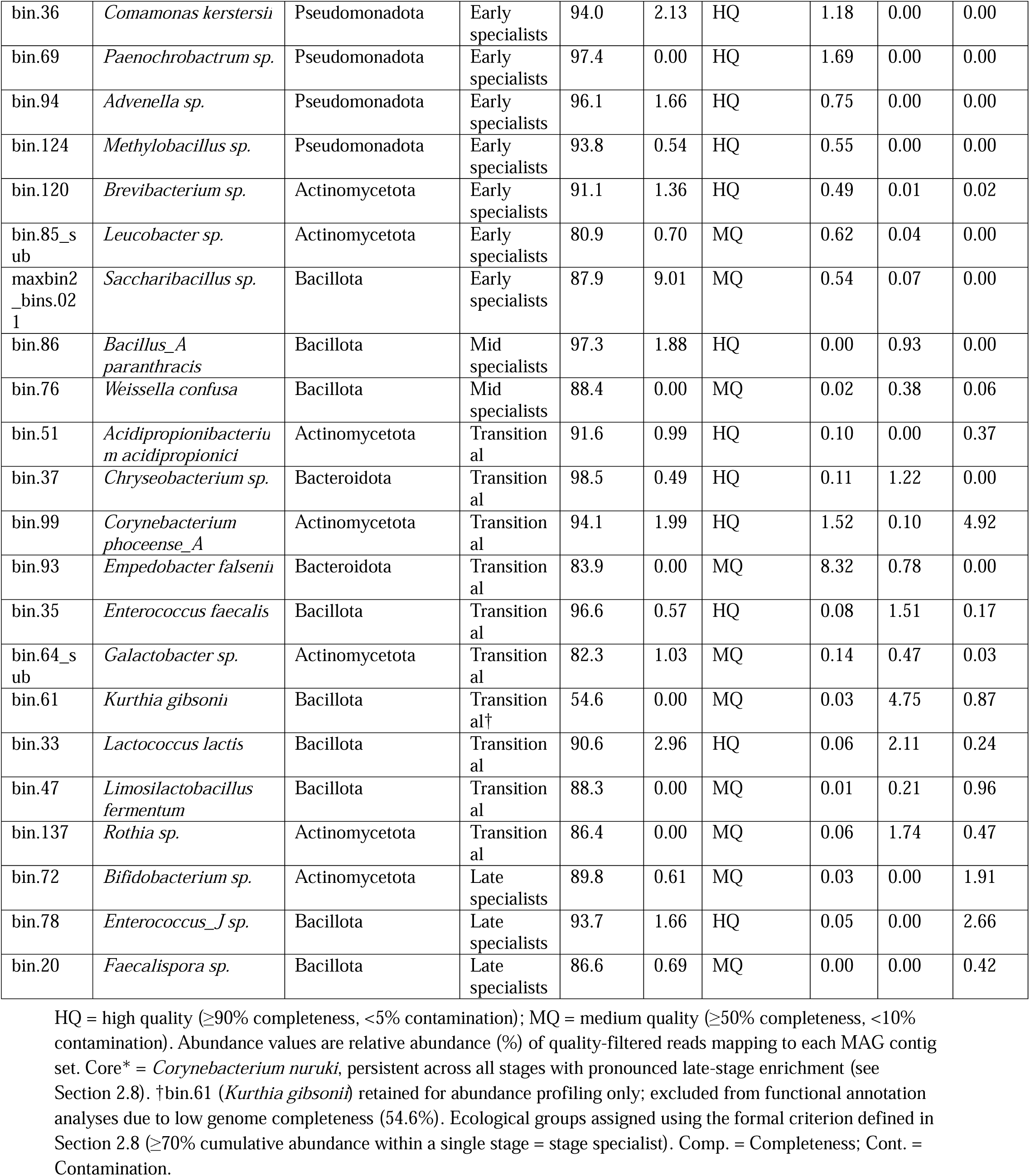
Ecological classification and stage-associated abundance patterns of retained MAGs during ugba fermentation

**Supplementary Table S4.** BioSample accession numbers for the metagenome-assembled genomes (MAGs) recovered during spontaneous ugba fermentation.

| MAG ID | Taxonomic assignment | BioSample accession |
| --- | --- | --- |
| bin.10 | *Corynebacterium variabile* | SAMN61874710 |
| bin.5 | *Corynebacterium nuruki* | SAMN61874711 |
| bin.20 | *Faecalispora* sp. | SAMN61874712 |
| bin.72 | *Bifidobacterium* sp. | SAMN61874713 |
| bin.78 | *Enterococcus_J* sp. | SAMN61874714 |
| bin.76 | *Weissella confusa* | SAMN61874715 |
| bin.86 | *Bacillus_A paranthracis* | SAMN61874716 |
| bin.137 | *Rothia* sp. | SAMN61874717 |
| bin.33 | *Lactococcus lactis* | SAMN61874718 |
| bin.35 | *Enterococcus faecalis* | SAMN61874719 |
| bin.37 | *Chryseobacterium* sp. | SAMN61874720 |
| bin.47 | *Limosilactobacillus fermentum* | SAMN61874721 |
| bin.51 | *Acidipropionibacterium acidipropionici* | SAMN61874722 |
| bin.61 | *Kurthia gibsonii* | SAMN61874723 |
| bin.64_sub | *Galactobacter* sp. | SAMN61874724 |
| bin.93 | *Empedobacter falsenii* | SAMN61874725 |
| bin.99 | *Corynebacterium phoceense_A* | SAMN61874726 |
| bin.119 | *Flavobacterium* sp. 1 | SAMN61874727 |
| bin.120 | *Brevibacterium* sp. | SAMN61874728 |
| bin.124 | *Methylobacillus* sp. | SAMN61874729 |
| bin.126 | *Flavobacterium* sp. 2 | SAMN61874730 |
| bin.146 | *Bacteroides* sp. | SAMN61874731 |
| bin.18 | *Sphingobacterium thalpophilum* | SAMN61874732 |
| bin.36 | *Comamonas kerstersii* | SAMN61874733 |
| bin.60 | *Dysgonomonas capnocytophagoides* | SAMN61874734 |
| bin.69 | *Paenochrobactrum* sp. | SAMN61874735 |
| bin.85_sub | *Leucobacter* sp. | SAMN61874736 |
| bin.89_sub | *Sphingobacterium* sp. 2 | SAMN61874737 |
| bin.94 | *Advenella* sp. | SAMN61874738 |
| maxbin2_bins.021 | *Saccharibacillus* sp. | SAMN61874739 |

